# Fluoxetine acts concomitantly on dorsal and ventral hippocampus to Trk-dependently modulate the extinction of fear memory

**DOI:** 10.1101/2021.01.28.428405

**Authors:** Cassiano Ricardo Alves Faria Diniz, Leandro Antero da Silva, Luana Barreto Domingos, Andreza Buzolin Sonego, Leonardo Resstel Barbosa Moraes, Sâmia Joca

## Abstract

**Background:** Hippocampus can be divided along its longitudinal axis into dorsal and ventral parts. Both are usually committed to modulate different aspects of behavior and stress response. However, it is not clear whether the hippocampal subregions could differently modulate the effect of antidepressant drugs. Since fluoxetine (FLX) effect on extinction of aversive memory is well known to depend on hippocampal BDNF levels, we hypothesized that the hippocampal subregions might play different roles in fluoxetine efficacy in decreasing fear response.

**Method:** Wistar rats were fear-cued conditioned and treated chronically with fluoxetine to enhance their subsequent extinction memory. First, FLX effect on BDNF levels was assessed considering the dorsal (dHC) and ventral (vHC) hippocampus apart. Then, K252a (a functional Trk blocker) was infused either into the dHC or vHC to assay its interaction with FLX treatment over the fear response. Next, BDNF was directly infused into either the dHC or vHC to compare its behavioral effects with FLX. Finally, FLX effect on c-Fos expression was evaluated also considering the dHC and vHC apart, along with subareas of amygdala and medial prefrontal cortex.

**Results:** Chronic FLX treatment increased BDNF in the dHC, whereas BDNF was increased in the vHC after acute treatment only. K252a infused after the extinction protocol into either dHC or vHC was able to prevent FLX effect on fear response. BDNF directly infused into the dHC increased fear response, however its administration into the vHC induced an opposite effect. Besides, a negative correlation between the fear response and c-Fos expression was observed after chronic FLX treatment specifically in the dHC CA3/CA1 and vHC CA1/DG.

**Conclusion:** Both dHC and vHC are important for the Trk-dependent FLX effect on extinction memory, although a discrepancy on the fear response was observed with the direct infusion of BDNF into the dHC or vHC.

## 1 INTRODUCTION

Intense trauma exposure may trigger a set of acute fear symptoms such as hyperarousal, avoidance and reexperiencing (Pai A, et al 2017). The majority of people usually extinguish those responses over time, however, the symptoms persist in some people to stablish the posttraumatic stress disorders – PTSD (Rothbaum BO and Davis M 2003). Traumatic experiences are common events, representing a lifetime risk as high as 50-60% of being exposed to any trauma, although a smaller portion 8-20% of the population develop in fact PTSD (Warner CH, et al 2013). Since PTSD represents a huge public health concern, it is important to search for a better comprehension of its neuropathology and possible new therapies.

PTSD is based on the strengthening of maladaptive associative memories inasmuch as neutral and harmless clues are previously experienced along with a traumatic event, then becoming the conditioned stimuli capable of inducing fear responses by themselves (Maren S and Holmes A 2016). Animal fear conditioning model has been used to mimic core PTSD characteristics, including the impaired extinction of conditioned responses. Extinction learning has also been stablished as an associative memory, in fact a new one which competes with the prior conditioned memory by retuning conditioned stimuli back to its neutral valence (Maren S and Holmes A 2016).

Accordingly, extinction memory is not permanent and conditioned responses are usually recovered by the reinstatement, renewal or spontaneous recovery of the fear symptoms (Quirk GJ and Mueller D 2008). Interestingly, animals treated for 14 days with fluoxetine (FLX), a selective serotonin-reuptake inhibitor (SSRI) used in PTSD treatment, presented enhanced extinction memory and avoided the re-collection of fear conditioned responses compared to non-treated animals (Karpova NN, et al 2011). However, FLX did not work as expected in BDNF (brain-derived neurotrophic factor) +/− heterozygous animals, thus indicating that BDNF is crucial to fluoxetine-induced effects (Karpova NN, et al 2011). BDNF usually activates tropomyosin receptor kinase B – TrkB (Soppet D, et al 1991), thus inducing TrkB autophosphorylation, to further induce neuroplasticity such as dendritogenesis, axonal sprouting, synaptic strengthening and neuronal survival (Park H and Poo MM 2013). In fact, antidepressants are suggested to work by boosting neuroplasticity through the reopening of the critical period, after which neurons usually decrease its capability for plastic flexibility (Castrén E and Antila H 2017).

However, BDNF effects on extinction memory and neuroplasticity seems to be rather complex. For instance, inducible BDNF knockdown in the dorsal hippocampus (dHC) weakened the extinction of fear-potentiated startle (Heldt SA, et al 2007), while reducing BDNF in the ventral hippocampus (vHC) impaired extinction of avoidance behavior (Rosas-Vidal LE, et al 2018). Moreover, BDNF infused directly into the vHC decreased fear conditioned response even without extinction training (Rosas-Vidal LE, et al 2014), whereas BDNF levels of the vHC was found increased after animal exposure to the avoidance (Rosas-Vidal LE, et al 2018) or cued fear (Peters J, et al 2010) extinction protocol. Through a translational perspective, hippocampal activity and its structural set up have also been shown to be modified in PTSD patients (Milad MR, et al 2009). Even though the whole hippocampus comprises a highly uniform intrinsic organization, its hodological characteristics and gene expression profile are quite different along the longitudinal axis (Strange BA, et al 2014; Fanselow MS and Dong HW 2010). Transcriptome and proteomic changes in response to stressful events, for example, have also shown different profiles along the dHC and vHC (Floriou-Servou A, et al 2018). From a behavioral overview, dHC functional role is described to be related to spatial navigation-based learning and memory, while vHC role has been associated to stress exposure-based behavioral and hormonal regulation (Bannerman DM, et al 2004).

Although antidepressant efficacy is usually associated with the regulation of hippocampal BDNF levels (Diniz CRAF, et al 2018), it is not totally clear whether dHC and vHC are differently involved. In order to figure out additional knowledge on this gap, BDNF role concerning the FLX effect on extinction memory has been searched in light of dHC or vHC.

## 2 METHODS

### 2.1 Animals

171 Male Wistar rats (250-350g) were housed in pairs in temperature-controlled room (24 ± 1ºC), with a 12-h light/12-h dark cycle (light on at 6:30 a.m.) and under standard laboratory conditions, including *ad libitum* access to food and water. *In vivo* experiments were conducted with approval of the local Ethical Committee (protocol 087/2015 for University of São Paulo), which are in accordance with the Brazilian Council for the Control of Animals under Experiment (CONCEA). All licenses comply with international laws and policies.

### 2.2 Drug Treatments

The following drugs were used: fluoxetine (FLX, selective serotonin reuptake inhibitor; Pharmanostra; 10mg/Kg i.p; (Karpova NN, et al 2011) diluted in saline 2% tween-80; K252a (inhibitor of Trk activity; Sigma-Aldrich, USA; 10pmol/500nl intra-hippocampal (Casarotto PC, et al 2015) diluted in saline 0,2% DMSO; BDNF (Sigma-Aldrich, USA; 0,25μg/0,05μl intra-hippocampal (Peters J, et al 2010) diluted in saline; Urethane (Sigma-Aldrich, USA, 5%, 10 ml/Kg i.p); Tribromoethanol (Sigma-Aldrich, USA, 2,5%, 10 ml/Kg i.p); Veterinary pentabiotic (1mL/Kg i.m) and anti-inflammatory Banamine (1mL/Kg s.c).

### 2.3 Surgery, Intracerebral Injections, and Histology

Rats were anesthetized with tribromoethanol to be fixed in a stereotaxic frame. Stainless steel guide cannulas (0.7mm OD) were implanted according to the rat brain atlas (Paxinos G and Watson C 1997) bilaterally into the dHC (AP= −4.0mm from bregma, L= +-2.8mm, DV= 2.1mm) or vHC (AP= −5.0mm from bregma, L= +-5.2mm, DV= 4.0mm). Cannulas were attached to skull bone with stainless steel screws and acrylic cement. A stylet inside the cannula was used to prevent any obstruction. Brain infusion was performed 5 to 7 days after surgery through a dental needle (0.3mm OD), in a total volume of 500nl/side per min (bilateral infusions). A Hamilton microsyringe (Sigma-Aldrich, USA) and infusion pump (KD Scientific, USA) were used for the infusions. Immediately after surgery, each animal received a dose of pentabiotic and anti-inflammatory to prevent postoperative infection and pain. In the case of hippocampal injections, rats were deeply anesthetized right after the end of the behavioral test with urethane, and 200nl of methylene blue was brain infused immediately prior to brain collection for site injection confirmation. Behavioral data corresponding to animals with injections outside the targeted area were discarded from statistical analysis. Representative sites of all brain infusions were inserted in diagrams (Figures 1a-d).

### 2.4 BDNF- ELISA

At the end of the behavioral experiments, animals were euthanized and dHC and vHC (ventral 2/3 of whole hippocampus) collected. Tissue was homogenized in a lysis buffer (20 mMTris-HCl pH=8; 137 mMNaCl; glicerol 10%) supplemented with protease inhibitor (Sigma-Aldrich, USA), and then centrifuged at 10.000g for 15min to the subsequent supernatant storage in −80°C. The content of BDNF was assessed by BDNF Emax EIA Assay System (Promega, USA), in accordance with manufacturer instructions. Bradford protein assay was used to quantify total protein levels (Bradford, Sigma-Aldrich, USA) and to normalize BDNF data.

**FIGURE 1.**
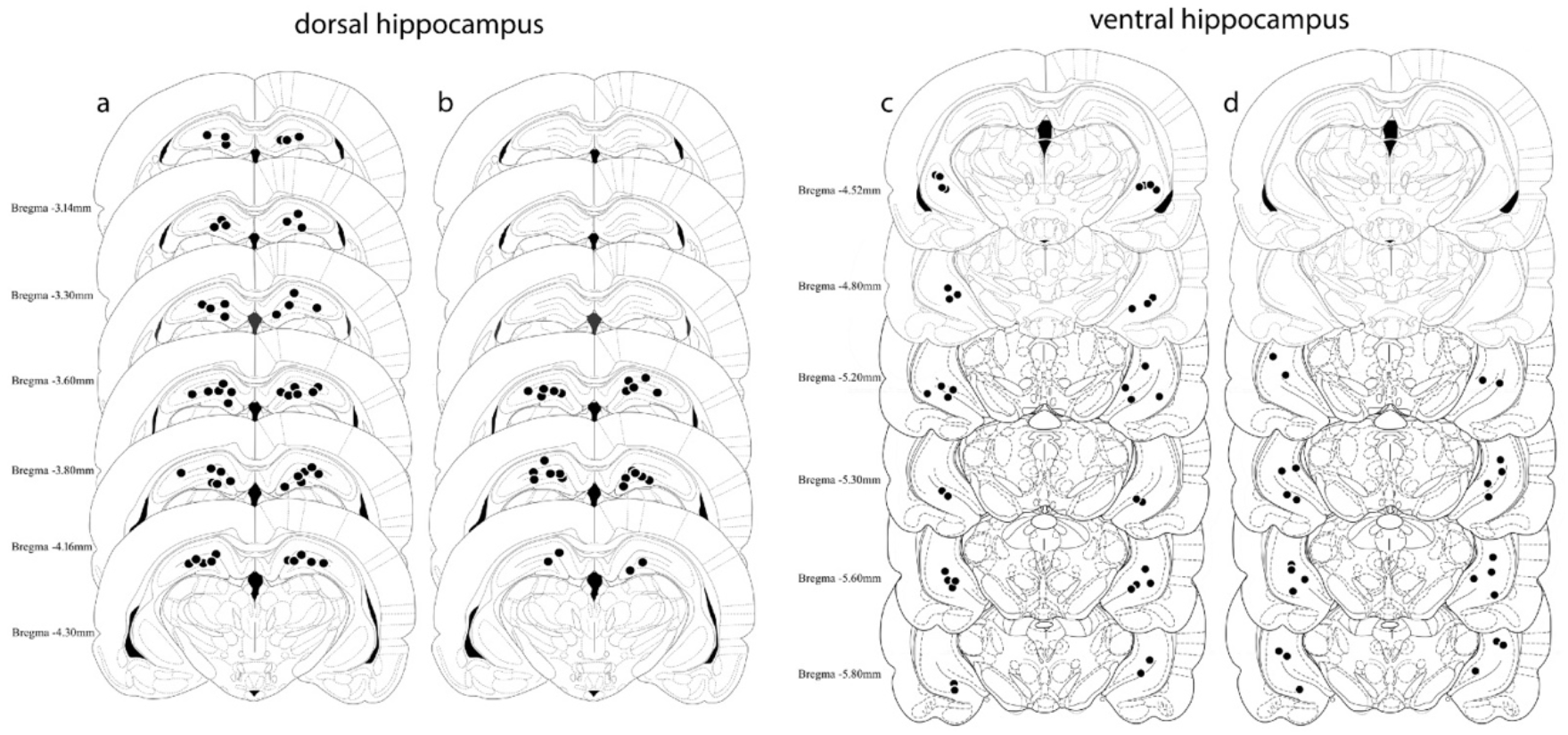
Histological localization of brain infusions. K252a (a and c) or BDNF (b and d) were infused into the dHC (left) or vHC (right) according to the pointed black circles diagramed on the atlas of Paxinos and Watson (1997). Due to overlap, the number of points represented is fewer than the real number of total infusions.

### 2.5 Hippocampal c-Fos protein immunohistochemistry

After a deep anesthesia, animals were submitted to a transcardiac perfusion with phosphate buffered saline (PBS) 0.01M followed by a 4% paraformaldehyde solution of 0.2M phosphate buffer (PFA 4%). Brains were collected and post-fixed overnight in PFA 4%, then immersed for 48h in a 30% sucrose solution of 0.01M PBS for cryoprotection. Further, brains were frozen in isopentane (−40°C) and sliced in serial coronal 40μm sections by using a cryostat (CM-1900, Leica, Germany). Samples were maintained in antifreeze solution before immunostaining procedure.

Free-floating sections were first rinsed (3 times/5min) in a 0.15% Triton-X solution of PBS 0.1M (pH 7.4, washing buffer) to further be pre-incubated for 30min with a 1% H_2_O_2_ solution of PBS 0.1M in order to avoid any confusing endogenous peroxidase activity. After that, samples had unspecific sites blocked for 1h in a 5% fetal bovine serum albumin solution of washing buffer to be next incubated overnight at 23°C with a polyclonal rabbit anti c-Fos antibody (1:1000 washing buffer, Santa Cruz Biotechnology, USA). After successive washing, slices were incubated for 1h in washing buffer containing a biotinylated goat anti-rabbit antibody (1:400; Vector Laboratories, USA). Next, sections were incubated with the avidin–biotin complex (1:200; Vector Laboratories, USA) for 2h. c-Fos immunoreactivity has been revealed by the addition of the chromogen 3,3′diaminobenzidine (DAB; 0.02%; Sigma–Aldrich, USA) in 0.1M PBS, to which 0.02% H_2_O_2_ was added before use. After ∼10 min, the tissue sections were rinsed, mounted on gelatine-coated slides and dehydrated.

c-Fos-positive cells were visualized under a 10× objective by using a light microscope (Olympus BX50, Japan) coupled to an Olympus camera (DP72, Japan). In the case of hippocampal sections, the number of c-Fos positive cells was counted manually by an observer blinded to the treatment scheme. About medial prefrontal cortex (mPFC) and amygdala, the software Image Pro-Plus 6.0 (Media Cybernetics, USA) counted as c-Fos positive the cells whose nuclei dark spot committed to an area between 10 and 80 μm^2^. Therefore, hippocampal cells were considered in raw numbers, and regarding mPFC and amygdala cell number is confined to a specific area (mm^2^). A total of two or three brain slices were used per animal. Since is not possible to differentiate left and right hemisphere, cells of the same slice were pooled together.

### 2.6 Cued fear conditioning

The apparatus consisted of two distinct environments, A for conditioning and B for all the other protocols (extinction, retention I and retention II). Context A was completely black with metal grids on floor, while context B was smaller, with a white and flat floor and black stripes along the white walls. All the apparatus was cleaned with 70% ethanol in between sessions. Animals were conditioned pairing 3 neutral sound stimuli (30s, White noise, 80dB) with an unconditioned stimulus (US, 1s foot shock 0,6mA, inter-trial interval: 20-120s). The US co-terminated with the neutral stimulus. Throughout the conditioning, the neutral stimulus became a conditioned one (CS).

Extinction protocol consisted of 21 CS (30s, White noise, 80dB, inter-trial interval: 20-60s). The first block of 3 CS was used as an average to set up the acquisition phase, while the last block of 3 CS was used as an average to set up the extinction phase. Next day, animals were exposed again to the B environment (short-term retention), but now only 6 CS were presented as prior described. Long-term retention was assessed 7 days after the short one, also with 6 CS presentation in environment B. Both retention protocols were set up based on the average of all 6 CS. Freezing has been defined as the percentage of time in which the animals are completely immobilized, except by breathing movements, during the 30s of sound stimulus. During each behavioral step animals were allowed to explore the environment for 2min without any stimulation. All the protocols were based on (Karpova NN, et al 2011), and evaluated by an experienced and treatment blinded observer. Of note, each behavioral experimental design was depicted along with the respective data graphics.

### 2.7 Statistical analysis

Data was analyzed with Student’s t test, or with one, two or three-way ANOVA, as appropriate. Post hoc has been done by using Tukey’s or Dunnett’s multiple comparisons test. Pearson’s correlation was performed to verify how c-Fos expression casually correlates with animal freezing. P value < 0.05 was considered to be significant. Data are shown as mean ± standard error of the mean (SEM). Graphpad Prism software was used to perform all the analysis.

## 3 RESULTS

In order to make our raw data free to be universally accessed, we provided all the prism and excel files available in repository figshare at (https://figshare.com/s/2b0904e80ddc4b211bae; DOI: 10.6084/m9.figshare.13564388).

### 3.1 Fluoxetine effect on BDNF levels in the dorsal and ventral hippocampus

First, we have confirmed that the chronic treatment with FLX (FLX cr), but not the acute treatment (FLX ac), is able to enhance fear extinction (figure 2a). Thus, twelve days of treatment with FLX enhanced extinction memory by decreasing fear response in short-term retention one day after the extinction protocol (One-way ANOVA F_2,32_=3.002, p=0.0638; Dunnetts post-hoc with adjusted pVEHxFLXcr=0.0373). Two-way ANOVA has found a time effect between acquisition and extinction (F_1,64_=24.41, p<0.0001), indicating a proper extinction procedure, however no effect of treatment (F_2,64_=0.8465, p>0.05) or any time x treatment interaction (F_2,64_=0.2262, p>0.05). Next, in an independent experiment, animals were conditioned (no difference of the raising fear response between groups – data not shown), treated as previously described, and euthanized one day after the last day of treatment to assay hippocampal BDNF levels. In order to focus on how FLX treatment change BDNF levels, we avoided any interference of the extinction protocol. The dHC BDNF levels have been increased only after chronic FLX treatment (One-way ANOVA F_2,20_=4.698, p=0.0213; Dunnetts post-hoc with adjusted pVEHxFLXcr=0.0218), while vHC BDNF levels was increased only after acute FLX treatment (One-way ANOVA F_2,20_=3.971, p=0.0353; Dunnetts post-hoc with adjusted pVEHxFLXac=0.0196) – figure 2b. These results suggest that the systemic FLX effect on extinction memory may depend differently on dHC and vHC-based BDNF change levels.

**FIGURE 2.**
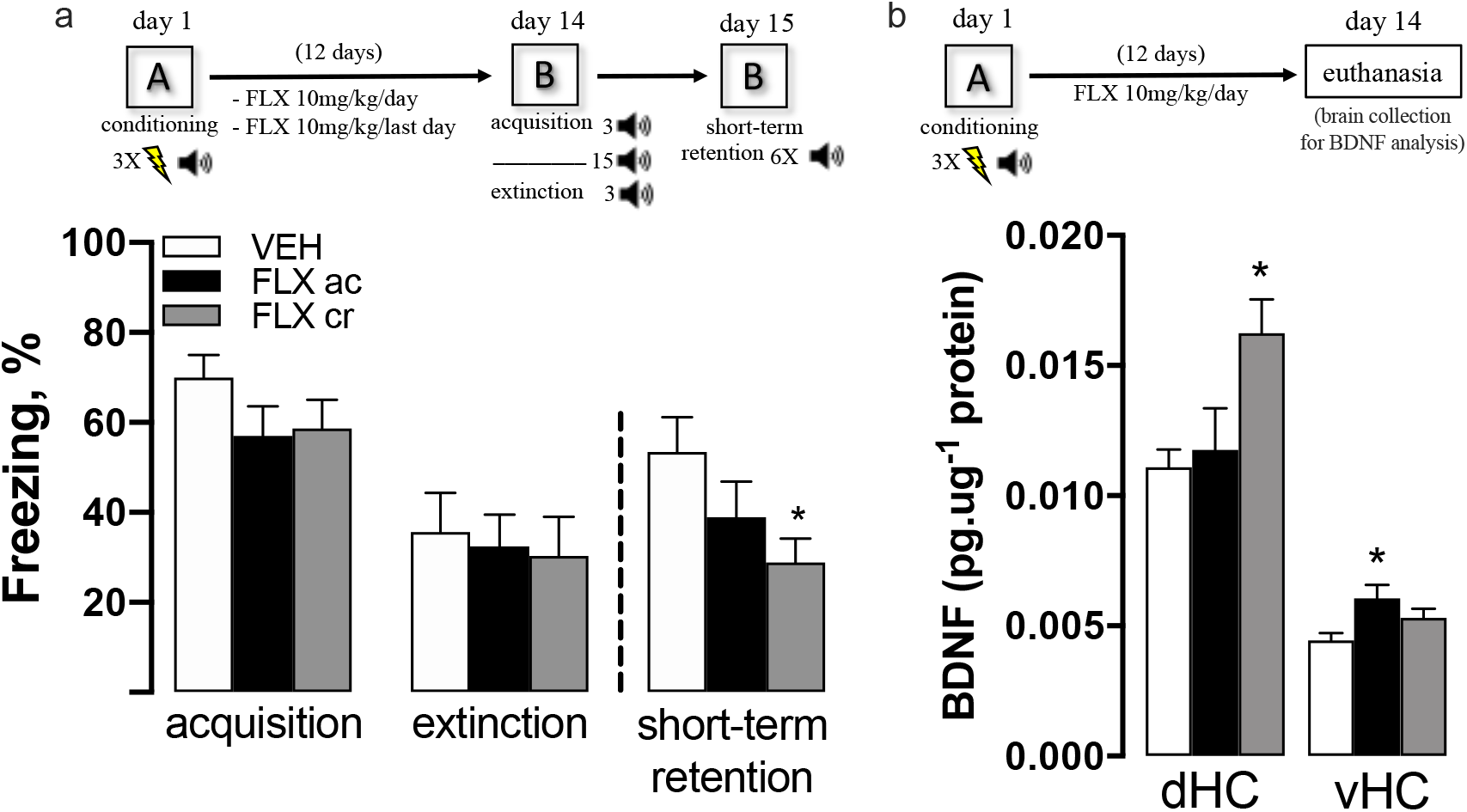
Fluoxetine effect on BDNF levels in the dorsal and ventral hippocampus. Animals were first submitted to the conditioning protocol (day 1), and then FLX (10mg/kg) or VEH treatment began one day after in a daily regime of i.p injection (twelve days). Acute group received VEH instead of FLX, except by the last day of treatment. (a) One day after the last day of treatment animals were through the extinction protocol (day 14), and short-term retention on day 15 (n: VEH 11, FLXac 12, FLXcr 12), (b) while other independent group of animals avoided extinction protocol to be directly euthanized on day 14 for BDNF brain measurements (n: VEH 7, FLXac 8, FLXcr 8). Data are expressed as mean ± SEM. *p<0.05 compared with VEH.

### 3.2 Fluoxetine effect on extinction memory depends on hippocampal Trk receptors

As described previously, after conditioning, the animals were treated with VEH or FLX chronically for twelve days. However, in this case the animals were submitted to the stereotaxic surgery six days before treatment with VEH or K252a either into dHC or vHC right after the extinction protocol. In both independent experiments, three-way ANOVA was capable of finding a time effect between acquisition and extinction (dHC: F_1,82_=40.43, p<0.0001; vHC: F_1,42_=82.68, p<0.0001), pointing for a proper extinction procedure. However, no effect of first (VEH/FLX) or second injections (VEH/K252a) has been found, neither any kind of interaction between factors (not shown). Regarding short-term retention, in dHC experiment the two-way ANOVA found an effect of first injections (VEH/FLX: F_1,41_=7.469, p=0.0092), no effect of second injections (VEH/K252a: F_1,41_=2.494, p=0.1219) and a trend in the interaction between the factors (F_1,41_=2.893, p=0.0966); while in vHC experiment no effect of first injections has been found (VEH/FLX: F_1,21_=1.636, p=0.2149), however the effect of second injections (VEH/K252a: F_1,21_=5.030, p=0.0358) and the interaction between factors (F_1,21_=6.580, p=0.0180) were significant. Still concerning short-term retention, the Tukey’s post-hoc was used and all the relevant adjusted p values are described in figure 3a and 3b. These results suggest that the systemic FLX effect on extinction memory may depend on the Trk found in both dHC and vHC.

**FIGURE 3.**
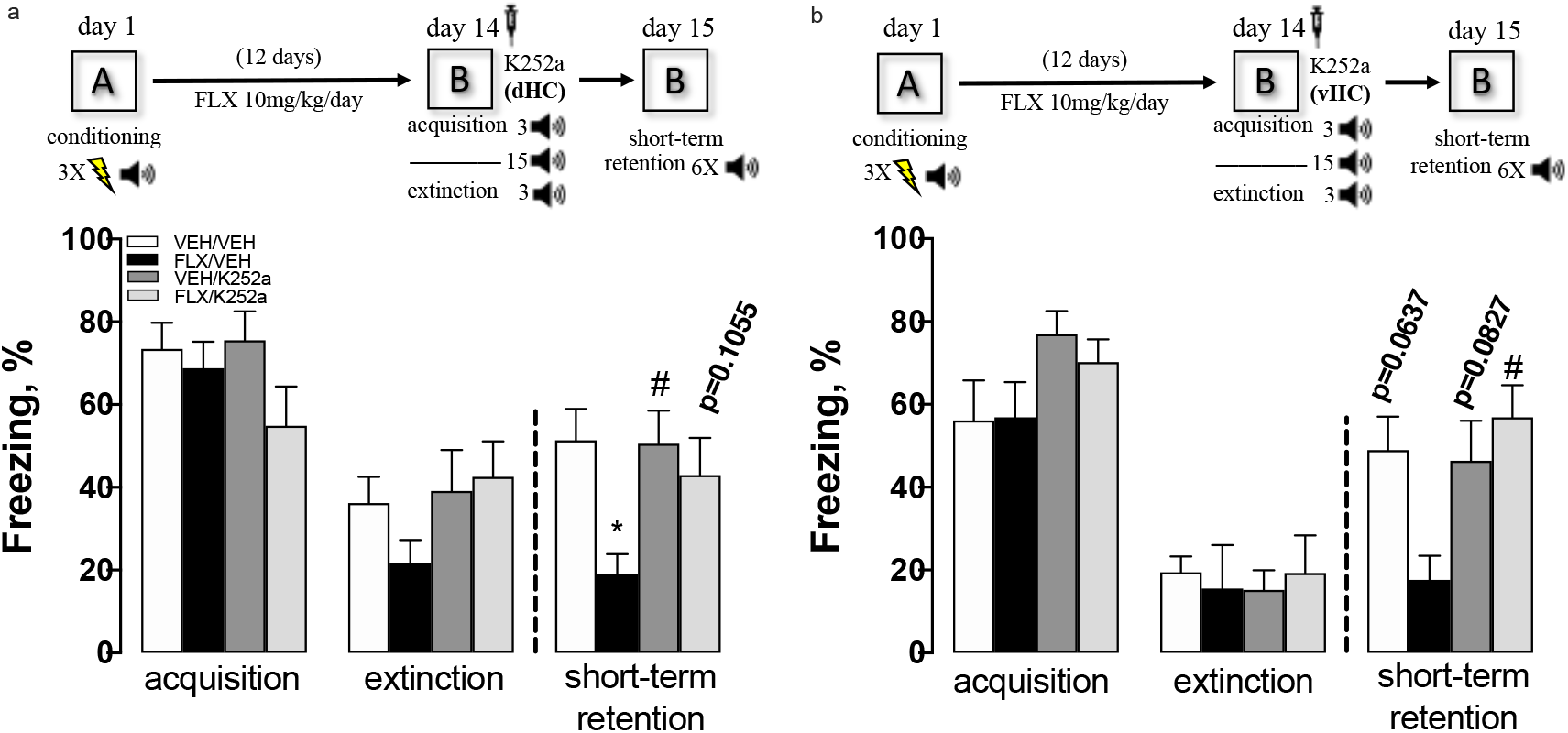
Fluoxetine effect on extinction memory depends on hippocampal Trk receptors. Animals were first submitted to the conditioning protocol (day 1), and then FLX (10mg/kg) or VEH treatment began one day after in a daily regime of i.p injection (twelve days). One day after the last day of treatment animals were through the extinction protocol (day 14), and short-term retention on day 15. Independent group of animals received K252a or VEH into (a) dHC (n: VEH/VEH 12, VEH/k252a 10, FLX/VEH 13, FLX/k252a 10) or into (b) vHC (n: VEH/VEH 6, VEH/k252a 7, FLX/VEH 6, FLX/k252a 6), right after the extinction protocol. Data are expressed as mean ± SEM. *p<0.05 compared with VEH/VEH. #p<0.05 compared to FLX/VEH. All p values described in the graph are set up by the comparison with FLX/VEH.

### 3.3 Effect of BDNF infused into dHV or vHC on extinction memory

After conditioning the animals were maintained undisturbed in their cages for twelve days up to extinction protocol, except by basic handlings and the mid-way stereotaxic surgery procedure. Independent batch of animals received BDNF into either dHC or vHC right after the extinction protocol, and two retention protocol was further stablished, the short and the long-term one. In both experiments, two-way ANOVA found a time effect between acquisition and extinction (dHC: F_1,24_=83.60, p<0.0001; vHC: F_1,22_=4.814, p=0.0391), pointing for a proper extinction procedure. However, in both cases no effect of treatment VEH/BDNF (dHC: F_1,24_=0.0169, p=0.8977; vHC: F_1,22_=0.1359, p=0.7160), neither an interaction between the factors (dHC: F_1,24_=1.511, p=0.2310; vHC: F_1,22_=0.0476, p=0.8294), was observed. Student’s t-test found that BDNF infusion into dHC increased fear response in short-term retention (t_12_=2.535, p=0.0262), but unchanged fear response in long-term retention (t_12_=0.0895, p=0.9302); while BDNF infusion into vHC unchanged fear response in short-term retention (t_11_=0.7397, p=0.4750), but decreased fear response in long-term retention (t_11_=2.945, p=0.0133) – figure 4a and 4b. Altogether, these results suggest that the bolus infusion of BDNF induces a dHC and vHC-based opposite action on extinction memory.

**FIGURE 4.**
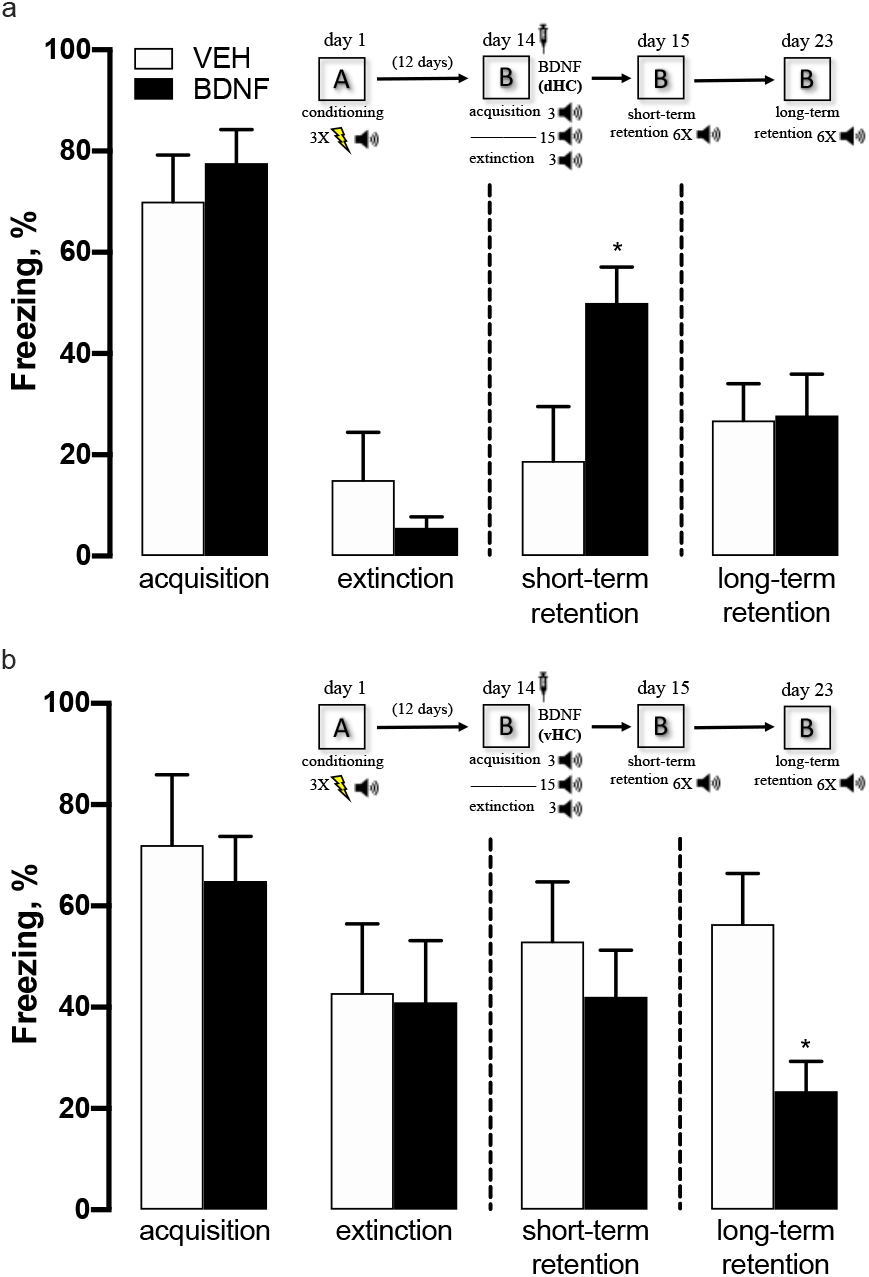
Effect of BDNF infused into dHV or vHC on extinction memory. Animals were first submitted to the conditioning protocol (day 1), and then to the extinction protocol on day 14. A short-term retention was provided one day after the extinction protocol (day 15), whilst the long-term retention was provided 7 days after the short-term one (day 22). Independent group of animals received BDNF or VEH either into (a) dHC (n: VEH 6, BDNF 8) or into (b) vHC (n: VEH 6, BDNF 7), right after the extinction protocol. Data are expressed as mean ± SEM. *p<0.05 compared with VEH.

### 3.4 Correlation of fluoxetine effect on extinction memory and c-Fos brain activity

Between conditioning and extinction protocols, animals were treated by twelve days either with FLX or VEH. A two-way ANOVA found a time effect between acquisition and extinction (F_1,26_=94.88, p<0.0001), pointing for a proper extinction procedure (figure 5a). An effect of treatment was also observed (F_1,26_=4.800, p=0.0376), however none interaction between the factors (F_1,26_=0.4933, p=0.4887). Student’s t-test found that FLX treatment decreased fear response in short-term retention (t_13_=2.297, p=0.0389) – figure 5a. One and a half hour after the end of previous behavioral procedure, animals were deeply anesthetized to be then transcardially perfused. Further, c-Fos immunostaining was developed as described in topic 2.5. Student’s t-test pointed that FLX treatment increased significantly the c-Fos expression in dHC CA3 (t_13_=2.360, p=0.0346), and a trend of increase was also observed in CA1 (t_13_=1.980, p=0.0692) and DG (t_13_=1.893, p=0.0808) – figure 5b. Otherwise, Student’s t-test showed a significant effect of FLX in increasing c-Fos expression all over the three vHC areas (CA3: t_13_=2.382, p=0.0332; CA1: t_13_=2.779, p=0.0156; DG: t_13_=3.129, p=0.0059) – figure 5b. FLX also increased c-Fos expression in the lateral amygdala (LA: t_13_=2.663, p=0.0195), but no effect was observed on basolateral (BA) or central (Ce) amygdala (BA: t_13_=1.469, p=0.1656; Ce: t_13_=1.732, p=0.1069) – figure 5c. No effect of FLX was found on ventral medial prefrontal cortex (vmPFC) areas (prelimbic – PL: t_13_=1.816, p=0.0924; infralimbic – IL: t_13_=0.6065, p=0.5546) – figure 5d. A Pearson’s correlation analysis between the animals' freezing level in short-term retention and the expression of c-Fos was performed over the respective brain areas where FLX influenced the most (p and Pearson’s r value available next to the graphs). In an overview, dHC CA3/CA1 and vHC CA1/DG are apparently the main areas whose FLX leverage over brain activity is likely to support extinction memory related-behavior change, figure 5e-g. An illustrative figure based on the atlas of Paxinos G and Watson C (1997) is shown with the bounded brain areas used for the c-Fos immunohistochemistry, figure 5h.

**FIGURE 5.**
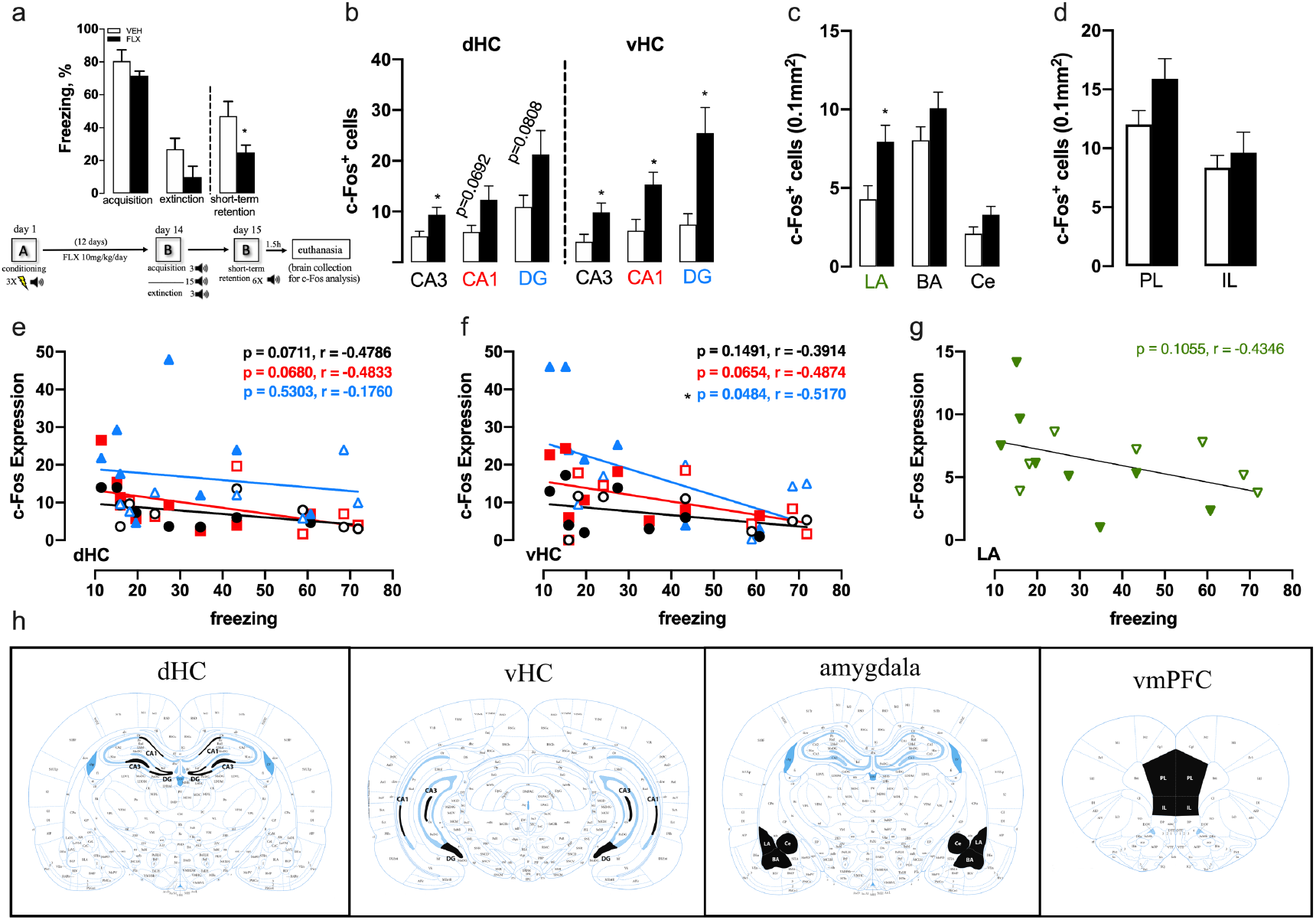
Correlation of fluoxetine effect on extinction memory and c-Fos brain activity. Animals were first submitted to the conditioning protocol (day 1), and then to the extinction protocol on day 14. A short-term retention was provided one day after the extinction protocol (day 15), and animals were euthanized one hour after the behavioral procedure to follow with c-Fos immunostaining procedures. Chronic FLX treatment (a) decreased fear response during short-term retention (n: VEH 7, FLX 8), increased (b) hippocampal and (c) LA c-Fos expression, while no change was observed in (d) vmPFC. (e,f,g) Pearson’s correlation between the animals' freezing level in short-term retention and the expression of brain c-Fos (filled geometries are FLX treated animals and the empty ones VEH treated). Atlas of Paxinos G and Watson C (1997) with the bounded brain areas (in black) used for the c-Fos immunohistochemistry (h). Data are expressed as mean ± SEM. *p<0.05 compared with VEH.

## 4 DISCUSSION

Fluoxetine has long been attributed to enhance extinction memory by reducing fear response after chronic treatment (Karpova NN, et al 2011; Popova D, et al 2014; Deschaux O, et al 2011; Deschaux O, et al 2012; Spennato G, et al 2008). Indeed, our fear conditioning protocol and drug treatment schedule corroborated it by showing reduced fear in the rats during the short-term retention protocol (fig 2a). The chronic treatment regimen also increased BDNF protein levels found in the dHC (fig 2b). Unexpectedly, an increment of BDNF protein levels was also observed after acute treatment, but only in the vHC (fig 2b). Although these data suggest that only changes in BDNF levels in the dHC would be important for triggering FLX behavioral effect, we next observed that K252a infused after extinction protocol into either the dHC or vHC prevented FLX effect (fig 3a and 3b). This, thus, indicates that Trk signaling in both dHC and vHC would be involved in FLX effects on extinction memory.

A recent work has shown that antidepressants, including fluoxetine, are actually able to directly bind to the TrkB to allosterically modulate BDNF action (Casarotto PC, et al 2020). So, it is reasonable to suggest that FLX along with basal levels of vHC BDNF are enough to induce a TrkB activity-dependent behavioral change, insofar as FLX is then supposed to qualitatively potentiate even the basal action of BDNF. Curiously, the increase of vHC BDNF protein levels after acute FLX treatment was not sufficient to induce a fast-behavioral change. In humans, antidepressants' concentration is accumulated along the continuous treatment up to reach a brain steady state of the drug concentration supposed to be effective (Bolo NR, et al 2000; Henry ME, et al 2000; Karson CN, et al 1993; Kornhuber J, et al 1995). In this sense, presumably the FLX brain concentration after acute treatment is still too small to boost any BDNF action on TrkB, even though vHC BDNF levels were already slightly increased. Remarkably, we were able to prevent FLX effect with only one infusion of K252a immediately after the extinction protocol. BDNF is sorted to be released through a constitutive (spontaneous release) or regulated (release in response to a stimulus) secretory pathway (Bai Lu, et al 2005). Therefore, it is tempting to suggest that the extinction engagement works as a stimulus to trigger BDNF release, then making K252a able to block FLX effect either dependent on the interaction with synaptically available basal (vHC) or increased (dHC) BDNF levels.

Further, BDNF was bolus infused immediately after the extinction protocol into either the dHC or vHC (fig 4a and 4b). BDNF mimicked the action of FLX to reduce the fear response only when infused into the vHC, although we observed this BDNF effect only along the long-term retention. Meanwhile, dHC BDNF infusion actually induced an unanticipated and apparently contradictory behavioral effect by expanding the fear response throughout the short-term retention test. In accordance with our results, BDNF acute infusion into the vHC was found to enhance the fear extinction memory of mice, thus reducing their fear response (Rosas-Vidal LE, et al 2014). Additionally, gradual or acute increase in BDNF concentrations has been found to evoke discrepant molecular signatures and synaptic rearrangements of the hippocampal cells (Ji Y, et al 2010). Therefore, behind our conflicting data it is likely that a gradual increase of the BDNF levels within the dHC after chronic FLX treatment may elicit different behavioral consequences than that observed with a bolus BDNF injection. Also, it is noteworthy that the effect of FLX is likely more complex than BDNF since FLX is known to act through several direct and indirect mechanisms, including the triggering of BDNF/TrkB triggering. Thus, any direct comparison between them must be taken parsimoniously.

Finally, we replicated the results of chronic treatment with FLX on the short-term retention test (fig 5a) to then euthanize the animals for developing c-Fos immunohistochemistry. Conditioned fear response of animals as well as posttraumatic stress disorder itself is well known to be regulated by the triad hippocampus, amygdala and prefrontal cortex (Zelikowsky M, et al 2014; Sierra-Mercado D, et al 2011; Orsini CA, et al 2011; Milad MR, et al 2009; Pitman RK, et al 2012), so with c-Fos approach we may imply the functional role of specific sub-regions regarding all of those different brain areas in order to detail the FLX effect simultaneously through a micro and macro (network) perspective. All sub-hippocampal regions such as CA3, CA1 and DG of both dHC and vHC had their number of c-Fos^+^ cells increased in FLX-treated rats (fig 5b). A negative correlation between c-Fos and freezing was confirmed only with dHC CA3/CA1 and vHC CA1/DG. Although the FLX effect of increasing cellular activity in dHC DG and vHC CA3 apparently is not directly correlated with the decrease of fear response, it is possible that the noisy and turbulent activity (decoherence) of these sub-areas is important to ignite and/or to flow toward the following sub-hippocampal area a coherent activity in order to settle an associative memory engram. Indeed, the hippocampal circuitry sets an organized and reverberating feed forward structure, and according to it BDNF directly infused into the DG is able to trigger an increase of both BDNF and c-Fos levels into the remaining CA3 and CA1 sub-hippocampal areas (Shirayama Y, et al 2002). The number of c-Fos^+^ cells was also found increased in LA, but not in BA or Ce, although this increment does not seem to be associated with the FLX-induced decay of the conditioned fear response (fig 5c and 5g). Additionally, PL and IL activity have been widely showed to modulate fear response, with PL and IL activity inducing, respectively, an strengthen or decline of the fear response. However, our data show no change at the number of c-Fos^+^ cells either in PL or IL area of the mPFC with FLX chronic treatment (fig 5d). Of note, the unchanged brain areas may still have specific sort of neurons to be activated or deactivated by FLX in order to influence the fear response. Altogether, in our conditions c-Fos data suggest the dHC CA3/CA1 and vHC CA1/DG as one of the most important brain areas to have their activity regulated by FLX to prompt the dampening of the fear response, thus endorsing the first experiments that depict both dHC and vHC as essentials for the Trk-dependent FLX effect.

Both dHC and vHC have been constantly described to modulate different categories of behavior and/or physiological response (Bannerman DM, et al 2004), however it is difficult to really pinpoint a qualitative clear functional distinction between them. For example, enhancement of dHC excitability induced by knockdown of HCN1 channels induced an anxiolytic-like effect in mice (Kim CS, et al 2012), even though the modulation of this phenotype has been previously linked to the tuning of the vHC (Bannerman DM, et al 2004). Therefore, it is worth to consider that actually these both areas may actually present superimposed function if you consider all the distinct techniques already addressed for inducing their respective perturbation. Also, dHC and vHC send indirect projections to each other (Fanselow MS and Dong HW 2010), and based on this communication between them the initial increment of BDNF levels observed specifically in the vHC could even be important for priming the subsequent FLX effect on BDNF levels of the dHC.

Concluding, our data show a mix of similar and distinct action regarding the hippocampal sub-areas (dHC and vHC) depending on the protocol approached, but in general lines we could state that these two brain areas are concomitantly important for modulating the Trk-dependent FLX effect on extinction memory even though they showed a different outcome to the direct BDNF infusion.

## ACKNOWLEDGEMENTS

The authors thank to Flavia F. Salata and Laura H. A Camargo for technical support. This work was supported by research grants from FAPESP (2018/18500-3 and 2018/04250-5) and CNPq.

## AUTHOR CONTRIBUTIONS

CRAFD, SRLJ and LBMR designed the study. CRAFD and LAS performed and analyzed the behavioral experiments. CRAFD, LBD and ABS performed the c-Fos immunostaining and analyzed it. CRAFD wrote the manuscript and all the authors reviewed it.

